# Coupling of Rho family GTPases during mesenchymal-to-epithelial-like transitions

**DOI:** 10.1101/244343

**Authors:** Christopher P. Toret, Pruthvi C. Shivakumar, Pierre-françois Lenne, Andre Le Bivic

**Affiliations:** Aix-Marseille Univ, CNRS, IBDM, Case 907, 13288 Marseille, Cedex 09, France

## Abstract

Many metazoan developmental processes require cells to transition between migratory mesenchymal- and adherent epithelial-like states. These transitions require Rho GTPase-mediated actin rearrangements downstream of integrin and cadherin pathways. A regulatory toolbox of GEF and GAP proteins precisely coordinates Rho protein activities, yet defining the involvement of specific regulators within a cellular context remains a challenge due to overlapping and coupled activities. Here we demonstrate that *Drosophila* dorsal closure is a simple, powerful model for Rho GTPase regulation during leading edge to cadherin contact transitions. During these transitions a Rac GEF elmo-dock complex regulates both lamellipodia and Rho1-dependent, actomyosin-mediated tension at initial cadherin contacts. Moreover, the *Drosophila* Rho GAP arhgap21 ortholog controls Rac and Rho GTPases during the same processes and genetically regulates the elmo-dock complex. This study presents a fresh framework to understand the inter-relationship between GEF and GAP proteins that tether Rac and Rho cycles during developmental processes.

## INTRODUCTION

Mesenchymal-to-epithelial transitions (MET) and epithelial-to-mesenchymal transitions (EMT) occur when cells change between migratory mesenchymal states and cell-cell adherent epithelial states. MET and EMT processes play central roles in metazoan development and disease, and are essential for gastrulation, neural crest, heart valve formation, palatogenesis, myogenesis, tumor metastasis and wound healing (Baum et al., 2008; Le Bras et al., 2012; Nieto, 2011; Thiery et al., 2009).

In MET-related processes, two opposing leading edges interact and establish a new cell-cell contact. Integrin and cadherin machineries are major regulators of MET-related processes and link to the versatile actin cytoskeleton(Baum et al., 2008; Harris and Tepass, 2010; Le Bras et al., 2012; Nieto, 2011). Actin filaments, whether assembled in Arp2/3-dependent branched structures or formin-dependent actomyosin cables, generate pushing or pulling forces within the cell (Goley and Welch, 2006; Murrell et al., 2015), Actin assembly pathways are largely regulated by Rho family GTPases, which are best characterized in mammalian cells (Burridge and Wennerberg, 2004). At a leading edge, downstream of integrin signaling, Rac activation promotes branched actin network formation in the lamellipodia for membrane extension, while in the lamellum, activation of RhoA results in the formation of contractile actin bundles or arcs that buttress, anchor and flatten the cell (Burnette et al., 2014; Guilluy et al., 2011). When adjacent leading edges interact, cadherins rapidly engage and actin is rearranged along the nascent cell-cell contact, which is coincident with a burst of Rac activity and diminished RhoA activity (Yamada and Nelson, 2007). Specific and localized regulation of Rho family GTPases is central to the precise control of critical actin-dependent processes during MET-like progression, but this regulation remains poorly understood.

A large family of guanine nucleotide exchange factors (GEFs) and GTPase activating proteins (GAPs) regulate the GTP-GDP state of Rho family proteins (Rossman et al., 2005; Tcherkezian and Lamarche-Vane, 2007). Ideally, unique GEFs and GAPs are thought to specify Rho family protein activity within different cellular regions and times, yet many GEFs and GAPs have promiscuous Rho protein specificities (Goicoechea et al., 2014; McCormack et al., 2013; Tcherkezian and Lamarche-Vane, 2007). In addition, several regulators are known to function in similar pathways, as illustrated by the multiple Rac GEFs that act at cadherin contacts (McCormack et al., 2013; Toret et al., 2014a), and confound the notion of uniqueness. Lastly, Rac and Rho activities are often inversely correlated in cells (Burridge and Wennerberg, 2004), which complicates interpretation of loss-of-function analyses as perturbations of one pathway likely affect the other.

The process of dorsal closure during *Drosophila* embryogenesis was previously described as a model for mammalian wound healing, but recent discoveries have highlighted mechanistic differences between the two processes (Belacortu and Paricio, 2011; Harden, 2002). During dorsal closure, two opposing epithelial layers, with a leading edge enriched with filopodia and lamellipodia occur, but they are primarily brought together by the contraction and apoptosis of the underlying embryonic amnioserosa cells (Gorfinkiel et al., 2009; Toyama et al., 2008). A contractile actomyosin band spans the leading edge of the epidermal cells from anterior to posterior canthi and maintains a taut cell front (Ducuing and Vincent, 2016; Pasakarnis et al., 2016; Solon et al., 2009). As the epithelial layers come together, filopodia at the leading edges align segments and zipper the cells together (Millard and Martin, 2008). Finally, new cadherin contacts sequentially form between the adjacent cells that seal the dorsal side of the embryo (Eltsov et al., 2015). These late events are poorly understood, but the transition from a leading edge to a cadherin contact make dorsal closure a promising model to study metazoan MET-like processes in a less complex genetic background and during an *in vivo* developmental process.

Rho family GTPases play critical roles during *Drosophila* dorsal closure (Harden et al., 1999), but the GEFs and GAPs that regulate epidermis leading edge dynamics and cell-cell contact formation remain poorly understood. A dorsal closure defect was originally described for *dock* mutants, an established Rac GEF that is in complex with elmo protein (Nolan et al., 1998). Based on its known role in mammalian integrin-mediated cell migration, the elmo-dock complex is thought to drive epidermis migration (Harden, 2002). However, an elmo-dock complex in mammalian cells was found recently to regulate Rho family GTPases transiently downstream of initial cadherin contact formation (Erasmus et al., 2015; Toret et al., 2014a). With roles downstream of both integrins and cadherins, the elmo-dock complex is in a prime position to regulate Rho family GTPases during MET. Therefore, we investigated the specific roles of the elmo-dock complex and a novel GAP protein on the MET-related Rac and Rho processes that occur during *Drosophila* dorsal closure.

## RESULTS

### Elmo-dock complex functions during cadherin-contact formation in *Drosophila*

To assess elmo-dock complex function during dorsal closure, stage 14-15 embryos that express endogenous DE-cadherin-GFP were imaged over 2 hrs in wildtype (wt), *dock, elmo*, and *elmo dock* mutants. The early steps of dorsal closure in elmo-dock complex mutants were indistinguishable from wt (Figure 1A and Supplemental movie S1 and S2), which suggests that early events such as potential epidermis migration pathways were unperturbed. However, at a late stage a terminal epidermal gap was observed in virtually all *elmo* and *dock* mutants (Figure 1A, arrowhead). In *elmo dock* mutants an additional enlargement of the posterior region prevented observation of a terminal gap (Supplemental movie S2). Upon dorsal closure completion in wt embryos, a DE-cadherin-labeled seam is observed along the length of the closure (Figure 1B). In *elmo* and *dock* mutants, a very late delay in closure that persisted for 15 minutes to over 2 hrs generated the terminal gap. This aberrant terminal gap was surrounded by irregular-shaped epidermal cells, which were separated by fragmented cellular structures and failed to establish DE-cadherin contacts (Figure 1B).

**Figure 1.**
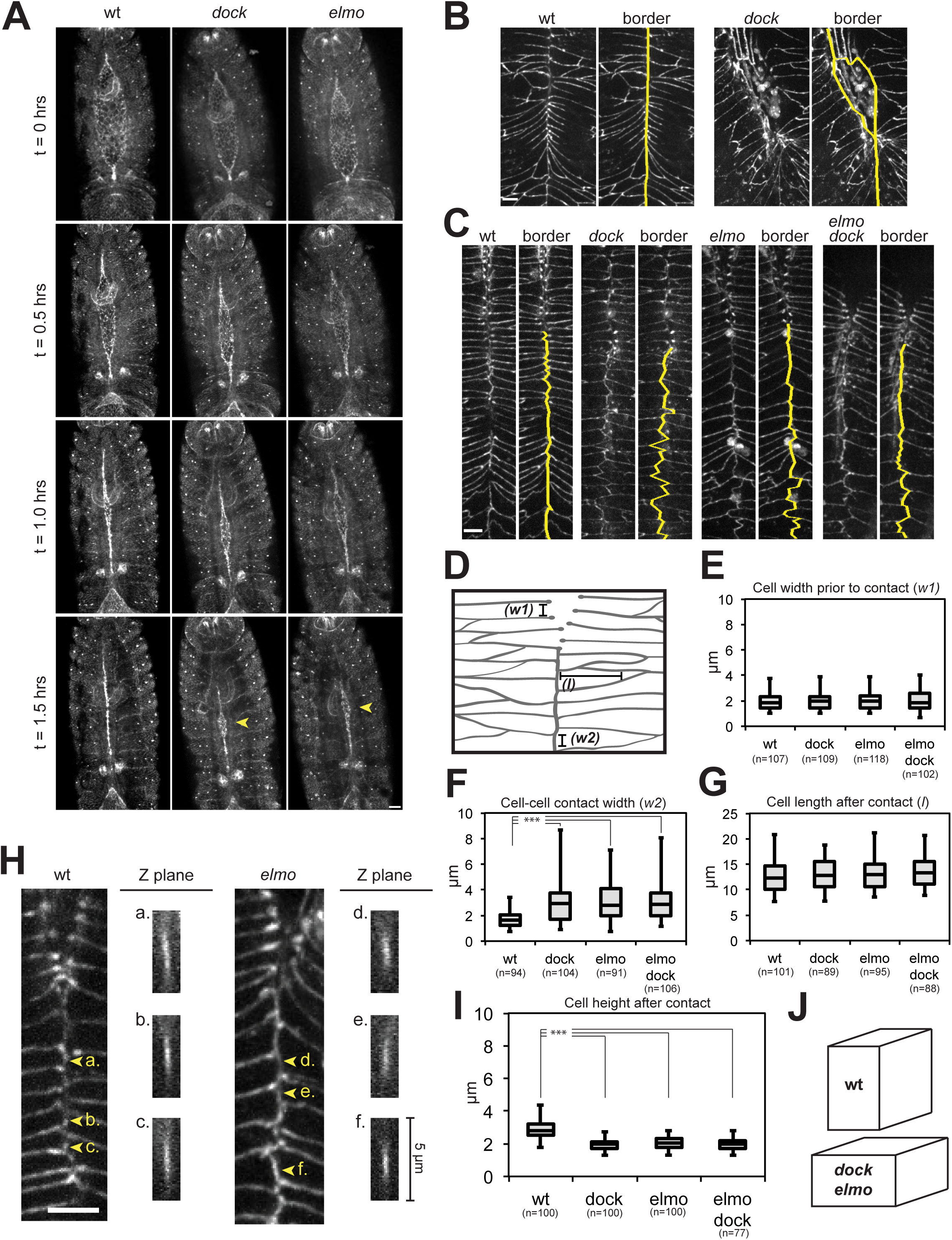
Elmo-Dock complex functions at new cell-cell contacts during dorsal closure. (A) Maximal intensity Z-stack projection of the dorsal side of wt, *elmo* and *dock* mutant embryos that express endogenous DE-cadherin-GFP over time. Yellow arrowheads indicate terminal gap. Scale bar = 25 μm. (B) Maximal intensity Z-stack projection of endogenous DE-cadherin-GFP expression during terminal phase of dorsal closure of wt and *dock* mutant (2 hrs after terminal gap appearance). Yellow lines indicate epidermis seams. Scale bar = 5 μm. (C) Maximal intensity Z-stack projection of endogenous DE-cadherin-GFP at cell-cell contact seams of dorsal closure in wt, *elmo, dock* and *dock elmo* mutants. Yellow lines indicate cadherin contact seams. Scale bar = 5 μm. (D) Schematic that indicates cellular dimensions measured in the zone of epidermal cadherin contact formation of dorsal closure. (E, F, G) Box plot of measurements of indicated cell dimensions for wt and mutant cells. Measurements pooled from 5-8 embryos and include anterior and posterior seams. (H) Maximal intensity Z-stack (left) with Z-sections of indicated cadherin contacts (right) in wt and *elmo* mutants. Scale bar = 5 μm. (I) Box plot of cell height measurements for wt and mutant cells. Measurements pooled from 6-8 embryos and include anterior and posterior seams. (J) Schematic representation of average cell shape of wt and mutants upon cell-cell contact formation.

To analyze cadherin-specific functions of the elmo-dock complex, DE-cadherin contacts along the closure seam were imaged in more detail. In wt embryos prior to cell-cell contact, DE-cadherin was strongly localized to cell contact edges near the leading edge (Figure 1C), as previously reported (Eltsov et al., 2015). Upon cell-cell contact formation, DE-cadherin was redistributed along the new cell-cell border, and no changes in overall cell widths before and after contact formation were observed (Figure 1C). In *elmo, dock*, and *elmo dock* mutants prior to cell-cell contact, DE-cadherin was distributed along the newly forming contacts similar to the wt, which suggests that the elmo-dock complex is not essential for this initial process (Figure 1C). However, upon cell-cell contact formation in these mutants, new DE-cadherin contacts appeared to expand along the anterior-posterior axis over a region of ~10 cells (Figure 1C). Thereafter this ~10 cell region older cell-cell contacts along the seam became irregular. (Figure 1C).

To quantify these defects, cell widths were measured for the 10 leading edge cells that preceded DE-cadherin contact formation (w1) and the 10 cells that formed new cadherin contacts (w2) (Figure 1D). The w1 measurements were found to be indistinguishable between wt and all of these mutants (Figure 1E). Analysis of new cadherin contacts (w2) revealed an expansion in *elmo, dock*, and *elmo dock* mutants (~3 μm) compared to wt (~ 2μm) (Figure 1F). Despite this expansion, no change in the lengths (l) of the cells after cell-cell contact formation was detected, which indicates that the mutant cells were either larger or flatter (Figure 1D and G). Z-stack images revealed differences in cell height in the 10 newest contacts between wt and mutants; wt cells had an average height of ~3 μm, while the mutants were ~2 μm (Figure 1H and I). Thus, wt cells appeared to extend in the Z-direction upon DE-cadherin contact formation, while mutant cells expanded along the axis of the closure (Figure 1J). These data indicate that elmo-dock complex activity is required immediately following DE-cadherin contact formation to increase the vertical height of new cell-cell contacts. Similar results with *dock, elmo*, and *elmo dock* mutants indicates that loss-of-function of either subunit is sufficient to block activity of the complex at new DE-cadherin contacts.

### The elmo-dock complex regulates DE-cadherin contact tension

Actomyosin-generated tension plays a major role in regulating dorsal closure on a tissue level (Solon et al., 2009; Wells et al., 2014). At a cellular level, actomyosin tension has an established role at cadherin contacts (Jean-Léon Maître, 2011). To test whether differences in cellular tension between wt and elmo-dock complex mutants were involved in cell shape changes, we used laser ablation of individual cell-cell contacts during dorsal closure. The initial velocity of adjacent cell border retraction is proportional to the tension of the ablated cell-cell contact with the assumption that the viscoelastic properties around the ablation site are unchanged across samples (Hutson et al., 2003; Rauzi and Lenne, 2011). Contacts were severed either prior to contact formation (leading edge), at new cell-cell contacts (1-10 cells), or at old cell-cell contacts (>10 cells after newest cell-cell contact) (Figure 2A). In the wt, ablation at the leading edge of an epidermal cell resulted in the rapid retraction of DE-cadherin borders that is indicative of high tension (Figure 2B, C and Supplemental Movie S3), and in agreement with the presence of an actomyosin cable spanning the leading edge of the epidermis (Solon et al., 2009). Border retraction upon ablation at the leading edge of *elmo* mutants was indistinguishable from wt (Figure 2B, C and Supplemental Movie S3), which indicates that the elmo-dock complex does not regulate actomyosin tension at the leading edge. Ablation of newly formed DE-cadherin contacts (1-10 cells) in wt or *elmo* mutant embryos revealed striking differences. In wt, little or no retraction was observed (Figure 2B, C and Supplemental Movie S3), which indicates that cells were under almost no tension upon initial cadherin contact formation. However in *elmo* mutants, cell borders still retracted, albeit at a reduced rate compared to leading edge levels, indicating a persistent tension at the new cell-cell contacts (Figure 2B, C and Supplemental Movie S3). Ablation of old contacts (>10 cells) showed that retraction was restored in the wt, though less than leading edge levels (Figure 2B, C and Supplemental Movie S3). In contrast, in *elmo* mutants there was essentially no retraction at older contacts (Figure 2B, C and Supplemental Movie S3). These data reveal that in wt cells tension at mature cell-cell contacts is restored, but this process fails in *elmo-dock* mutants. This may be a result of residual effects from initial cadherin contact defects or an additional elmo-dock complex role at mature contacts, and was not explored further. Ablation at cells directly adjacent to the newest formed cadherin contact revealed that cellular tension levels dropped rapidly upon DE-cadherin accumulation in wt cells (Figure 2D). Together these data indicate that elmo-dock complexes play a major role in the rapid regulation of cell-cell contact tension in response to initial cadherin engagement.

**Figure 2.**
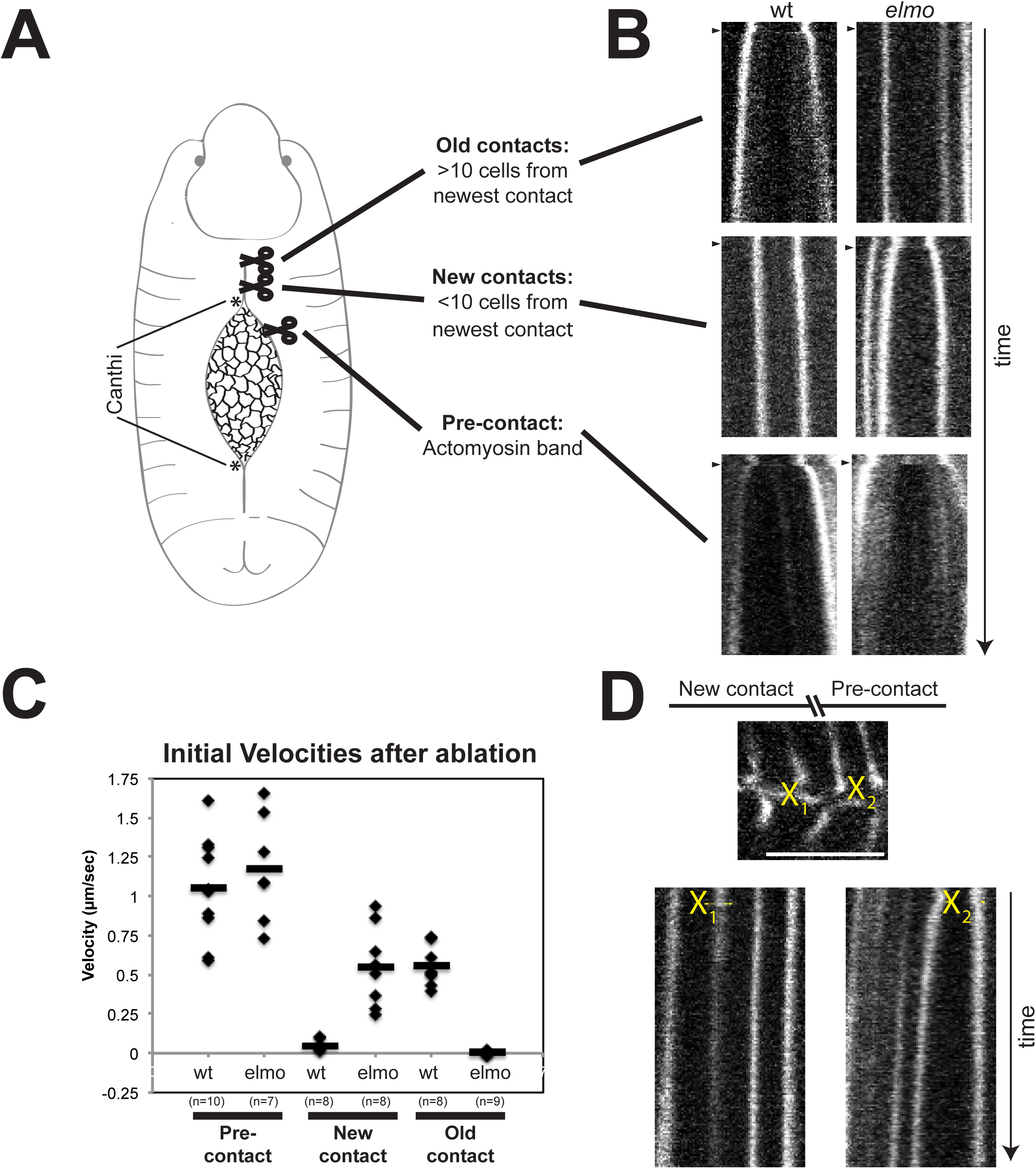
Epidermal cell-cell contact tension during dorsal closure. (A) Schematic, which depicts epidermis regions targeted for laser ablation. (B) Kymographs of cell-cell contact borders (DE-cadherin-GFP) flanking ablation site in indicated dorsal closure region. Arrowheads indicate moment of ablation. (C) Plot of initial velocity values calculated from cell vertex separation following ablation in indicated groups. Black bar indicates calculated average value. (D) Top: Still image from Supplemental Movie S4. Yellow ‘X’ indicates contact ablation sites. Scale bar = 5 μm. Bottom: Kymographs of cell-cell contact borders (DE-cadherin-GFP). Ablation site indicated with yellow ‘X’.

### The elmo-dock complex regulates myosin II and Rho1 at new DE-cadherin contacts

Nonmuscle myosin II and Rho1 regulate actomyosin-generated tension prior to cell-cell contact formation (Franke et al., 2005). To examine the localization of myosin II at DE-cadherin contacts during dorsal closure, mcherry-tagged myosin II regulatory light chain was imaged in conjunction with DE-cadherin-GFP in wt and *dock* mutants. Myosin II was enriched at the leading edges in wt, as reported previously (Franke et al., 2005). A similar localization was found at the leading edge of *dock* mutant cells (Figure 3A and B). Immediately upon cell-cell contact in wt cells, myosin II levels decreased at the newest DE-cadherin contacts, and the myosin II border progressed as new cell-cell contacts were formed (Figure 3A and B). In contrast, *dock* mutants retained myosin II along new cell-cell contacts, and the myosin II signal slowly tapered off after DE-cadherin contacts had formed (Figure 3A and B).

**Figure 3.**
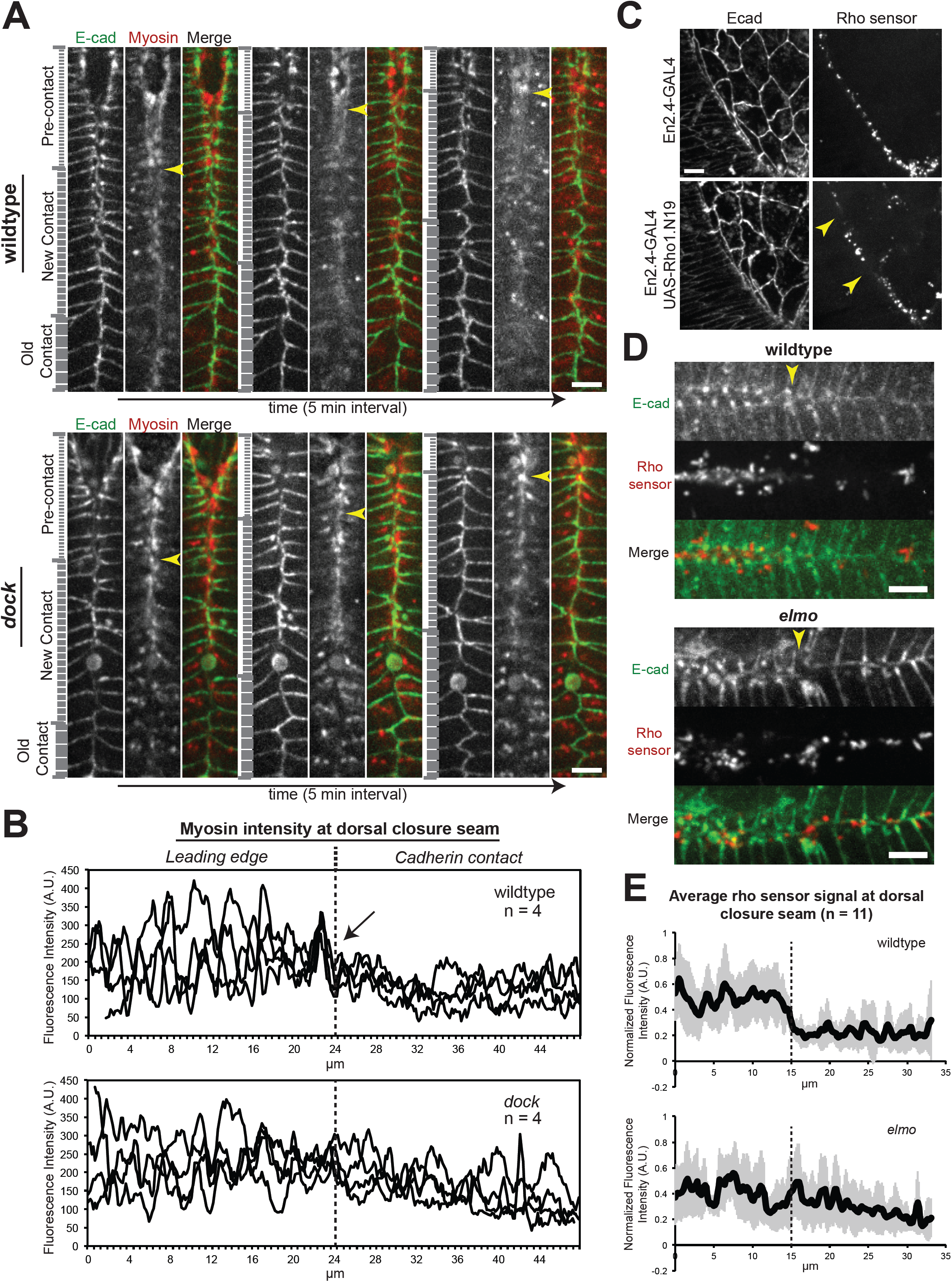
Nonmuscle myosin II and Rho1 GTPase activity in wt and *elmo-dock* mutants. (A) Maximal intensity Z-stack projections of wt and *dock* mutants, which express endogenous DE-cadherin-GFP and exogenous spaghetti squash-driven spaghetti squash-mCherry. Yellow arrowheads indicate most recent cell-cell contact. Scale bar = 5 μm. (B) Plots of background-subtracted fluorescence intensities measured along a 20 pixel-wide line that ran along and straddled one epidermal leading edge into cell-cell contact regions from 4 different embryos. Measurements were aligned to the newest cadherin. (C) Maximal intensity Z-stack projections of leading edge planes of wt and engrailed-GAL4; UAS-Rho1.N19 embryos that express endogenous DE-cadherin-Tomato and ubiquitin-driven Rho1 sensor-GFP. Arrows indicate regions lacking Rho sensor signal. Scale bar = 5 μm. (D) Maximal intensity Z-stack projections of wt and elmo mutants, which express endogenous DE-cadherin-Tomato and ubiquitin-driven Rho1 sensor-GFP. Yellow arrowheads indicate most recent cell-cell contact. Scale bar = 5 μm. (E) Plots of average fluorescence intensities with standard deviations calculated from mean intensities measured from 20 pixel-wide line that ran along and straddled one epidermal leading edge into cell-cell contact regions. Measurements were normalized and aligned to the newest cadherin contact before averaging.

Since Rho1-GTP is an activator of nonmuscle myosin II (Munjal et al., 2015), we examined Rho1 activity during dorsal closure. The anillin Rho-binding domain fused to GFP binds to Rho1-GTP acts as a biosensor for Rho1 activity (Munjal et al., 2015). During dorsal closure the Rho sensor formed puncta along the dorsal epidermal leading edges, which was lost segmentally upon engrailed-Gal4 expression (en2.4-GAL4) of a dominant negative Rho1 mutant, Rho1.N19 (Figure 3C). These results indicate that the biosensor puncta reported Rho1 activity in the epidermal cells of dorsal closure.

The anillin Rho sensor was analyzed in wt and *elmo* mutants to determine the levels of Rho1 activity during DE-cadherin contact formation in dorsal closure. In both wt and *elmo* mutants, the Rho sensor was localized in puncta along the leading edge of the epidermal cells. However, there was a decrease in Rho sensor accumulation in wt cells upon DE-cadherin contact formation; in contrast Rho sensor accumulation continued in *elmo* mutants after cell-cell contact formation (Figure 3D). Quantification revealed a cumulative sharp drop in Rho sensor aggregation upon DE-cadherin contact formation in wt (Figure 3E), whereas the Rho sensor signal in *elmo* mutants at best showed a gradual tapering off (Figure 3E). These results indicate that the elmo-dock complex is required for the rapid inactivation of Rho1 and loss of myosin II activity at new DE-cadherin contacts, which is consistent with the decrease in tension upon DE-cadherin contact formation (Figure 2).

### *Drosophila* arhgap21 regulates dorsal closure

Loss of elmo-dock complex function, a Rac GEF, resulted in aberrant Rho1 activity at new DE-cadherin contacts, but it is unknown how Rac regulators also influence Rho signaling. Recent models suggest that scaffolding between GEFs and GAPs is critical to coordinate Rho family GTPase cycles (Duman et al., 2015). The original screen that identified a cadherin function for the elmo-dock complex also identified the poorly characterized Rho GAP protein, rhogap19d (Toret et al., 2014b). Rhogap19d is the *Drosophila* ortholog of mammalian arhgap21/23, and arhgap21 is demonstrated to preferentially activate RhoA and RhoC in cells (Barcellos et al., 2013; Lazarini et al., 2013). This activity makes the rhogap19d protein, henceforth called *Drosophila* arhgap21, a strong candidate to coordinate Rho1 activity with the elmo-dock complex.

A role for arhgap21 in dorsal closure was investigated by ectodermal expression (P{GawB}69B) of arhgap21 RNAi (P{UAS-RhoGAP19D-dsRNA}). UAS-Lifeact-RFP embryos that expressed endogenous DE-cadherin-GFP were imaged with or without arhgap21 depletion. Unexpectedly, arhgap21-depleted embryos completed dorsal closure faster than wt embryos (Figure 4A and Supplemental Movie S4). This difference was apparent in the epidermis (DE-cadherin-GFP), while reduction of the amnioserosa appeared largely unaffected (Lifeact-RFP) (Figure 4A and B). Analysis of the rates of epidermal leading edge closure revealed that wt embryos closed at a rate of ~1 μm/min, and arhgap21-depleted embryos closed at a rate of ~1.4 μm/min (Figure 4C). In contrast to the epidermis, the speed of amnioserosa reduction was indistinguishable between wt and arhgap21-depleted embryos (Figure 4D). Similar results were obtained with a truncated arhgap21 mutant, which demonstrates that RNAi arhgap21-depletion effects were due to a loss of arhgap21 function (Supplemental figure S1A and B), and a TRiP line RNAi construct, which further suggests that the phenotype was not due to off-target effects (Figure 4C and D). The amnioserosa cell extrusion rates over one hour in wt, *elmo*, and arhgap21-depleted embryos were 11.7±1.5 %, 12.3±2.5 % and 10.7±3.0 % (n = 3 embryos each), respectively. Amnioserosa cell actomyosin pulse durations were also similar in wt (2.23±0.76 min), *elmo* mutants (2.14 ±0.65 min) and arhgap21-depleted (1.95±0.71 min) cells (n = 23 cells from 3 embryos each). These results are similar to previous observations (Cormier et al., 2012; Pasakarnis et al., 2016; Solon et al., 2009; Toyama et al., 2008), and suggest that arhgap21 does not significantly affect aminoserosa reduction mechanisms.

**Figure 4.**
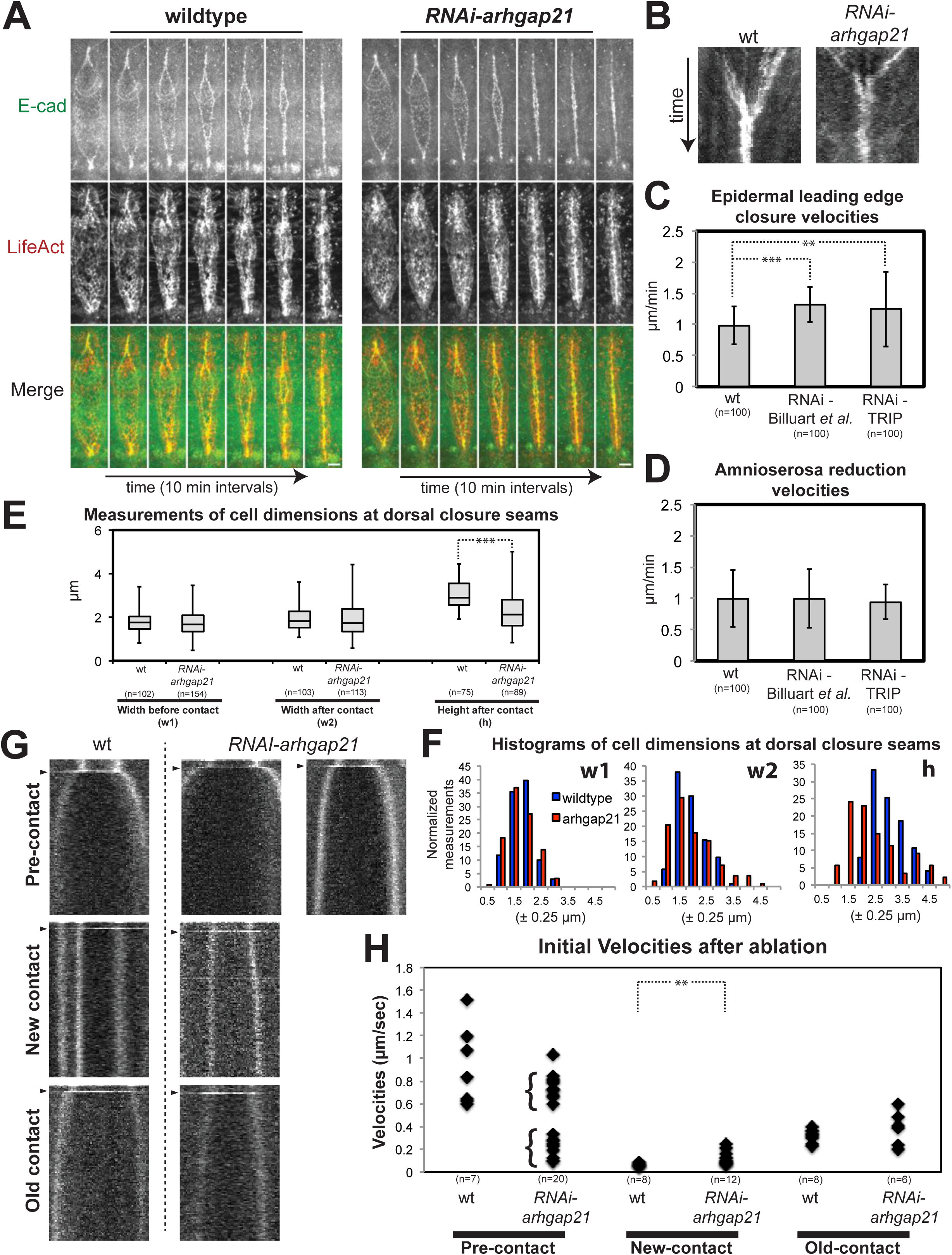
Arhgap21 functions in epidermal dorsal closure. (A) Montages of maximal intensity Z-stack projections of embryos, which express endogenous DE-cadherin-GFP and ectodermal Gal4-driven UAS-Lifeact-RFP with and without UAS-arhgap21-RNAi. Scale bar = 25 μm. (B) Kymographs of DE-cadherin-GFP during dorsal closure in wt and arhgap21-depleted embryos. Brightest signal indicates epidermal leading edges and converge at seams over time. (C) Quantifications of the change in linear distance that separate leading edges (DE-cadherin-GFP) during dorsal closure in 5 min intervals. Measurements pooled from three embryos from each of the indicated conditions. (D) Quantifications of the change in linear distance that spans the aminoserosa cells (LifeAct-RFP) during dorsal closure in 5 min intervals. Measurements pooled from three embryos from each of the indicated conditions. (E) Box plot of measurements of indicated cell dimension for wt and mutant cells. Measurements pooled from 5 embryos and include anterior and posterior seams. (F) Histograms of normalized cell dimensions for wt and mutant cells. (G) Kymographs of cell-cell contact borders (DE-cadherin-GFP) that flank the ablation site in indicated dorsal closure region (See Figure 2A). Arrowheads indicate moment of ablation. (H) Plot of initial velocity values calculated from cell vertex separation following ablation in indicated groups.

We also analyzed cell width and height during DE-cadherin contact formation in wt and arhgap21-depleted cells. No significant difference was detected between wt and arhgap21-depleted cell widths either before (Kolmogorov-Smirnov test for w1 is P=0.513) or after (Kolmogorov-Smirnov test for w2 is P=0.053) DE-cadherin contact formation (Figure 1D and Figure 4E). However, a significant difference in new cadherin contact heights was observed (Kolmogorov-Smirnov test for h is P<0.0005). Arhgap21-depleted contacts showed dual cell height peaks with a majority of shorter-than-wt cells, but an additional taller-than-wt cluster (Figure 4E and F). These results reveal an arhgap21 requirement for proper cell heightening after DE-cadherin contact formation.

Analysis of cell tensions in regions similar to those analyzed in the elmo-dock mutants (Figure 2A) revealed a significant difference in tension at the leading edge in wt and arhgap21-depleted cells prior to DE-cadherin-contact formation. In contrast to wt cells, arhgap21-depleted cells appeared to cluster in two populations of different tension, one of which was notably lower than the observed wt tension (Figure 4G and H). Tension levels in wt and mutants decreased upon new cadherin-contact formation (1-10 cell region), although not as significantly in arhgap21-depleted cells (Figure 4G and H). Lastly, no differences in tension were detected between wt and *arhgap21* RNAi at old contacts (>10 cells) (Figure 4G and H). In summary, this analysis revealed that arhgap21 regulates tension at both leading edge and new cell-cell contact during dorsal closure.

Embryos that express a UAS-LifeAct-GFP reporter with a Trojan GAL4 MiMIC insertion (Diao et al., 2015; Nagarkar-Jaiswal et al., 2015) were imaged, which allowed for the expression of GAL4 under the endogenous control of dock or arhgap21. This analysis revealed a ubiquitous expression for dock and arhgap21 during dorsal closure, although a strong labeling of actomyosin cables and seams was present (Supplemental figure S1A and C). We next tested whether there was a genetic interaction between arhgap21 and the elmo-dock complex. Embryos that ectodermally expressed arhgap21 RNAi in *elmo* mutants, and *arhgap21 dock* double mutants died prior to stage 14 (Supplemental figure S1D and E) indicating a synthetic lethal defect occurs prior to dorsal closure. Together, these data indicate that, like the elmo-dock complex, arhgap21 functions during dorsal epidermis closure. However, despite a synthetic defect, the consequence of a loss-of-function of the elmo-dock complex and arhgap21 are strikingly different.

### Dorsal closure actin structures in *elmo-dock* and *arhgap21* mutants

The loss-of-function phenotypes and known activities of the elmo-dock complex and arhgap21 suggest a crucial role for both proteins in the regulation of the actin cytoskeleton at the leading edge and cell-cell contacts of epidermal cells during dorsal closure. In order to visualize epidermal actin cables, LifeAct-RFP expressed in the ectoderm was imaged in wt, *elmo* mutant and arhgap21-depleted embryos. Actin cables were present as compact structures enriched in LifeAct-RFP signal at the leading edge of dorsal closures in wt and *elmo* mutant embryos (Figure 5A). In arhgap21-RNAi embryos, the cables appeared less defined with gaps in the LifeAct-RFP signal (Figure 5A).

**Figure 5.**
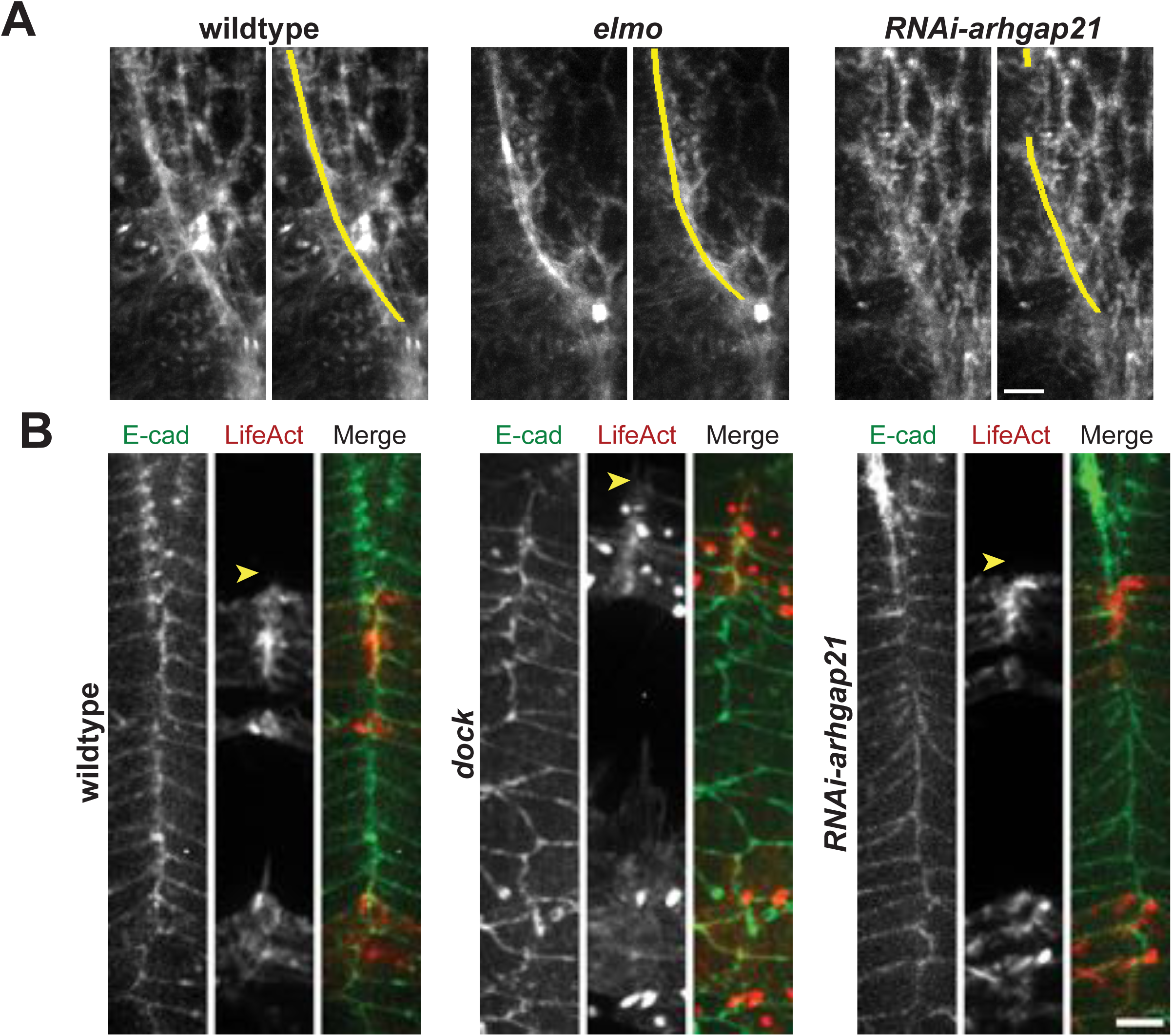
Dorsal closure actomyosin cable and cadherin contact actin structures. (A) For each indicated genetic background, (right panel) maximal intensity Z-stack projections of embryos, which express ectodermal Gal4-driven UAS-LifeAct-RFP, and (left panel) identical image with actomyosin band labeled in yellow. Scale bar = 25 μm. (B) For each indicated genetic background with engrailed-Gal4 UAS-LifeAct-GFP and endogenous DE-cadherin-Tomato, maximal intensity Z-stack projections of cell-cell contact seam. Arrowheads indicate most recent cell-cell contact. Bright red foci apparent in *dock* mutant are sites of actin-rich denticle initiation. Scale bar = 25 μm.

To analyze actin structures at epidermal cadherin contacts, UAS-LifeAct-GFP was expressed in engrailed stripes (en2.4-GAL4), which gave epidermal-specific expression. No differences in the overall distribution of LifeAct-GFP were detected between wt, *dock* mutant, and arhgap21-depleted new (1-10 cells) and old (>10 cell) contacts. For all conditions, new cell-cell contacts appeared enriched in LifeAct-GFP signal and diminished at older contacts (Figure 5B). However, LifeAct-GFP may not effectively distinguish between Rac- and Rho-dependent actin assemblies at cadherin contacts.

Engrailed-driven UAS-LifeAct-GFP distinguished different actin structures in the epidermis leading edge (filopodia, lamellipodia, and cable) (Figure 6A). Quantification of the number of filopodia structures present at leading edges revealed no significant difference between wt, *elmo* or arhgap21-depleted embryos (Figure 6B). However some differences occurred in the dynamics of filopodia (Supplemental Movie S4); tracking of filopodia tips revealed that the lateral probing behavior of filopodia in *dock* mutants was mildly reduced compared to wt, while filopodial movements were mildly enhanced in *arhgap21*-depleted cells relative to wt (Figure 6C). Effects on filopodia dynamics may be linked to Cdc42 pathways and were not pursued further in this study.

**Figure 6.**
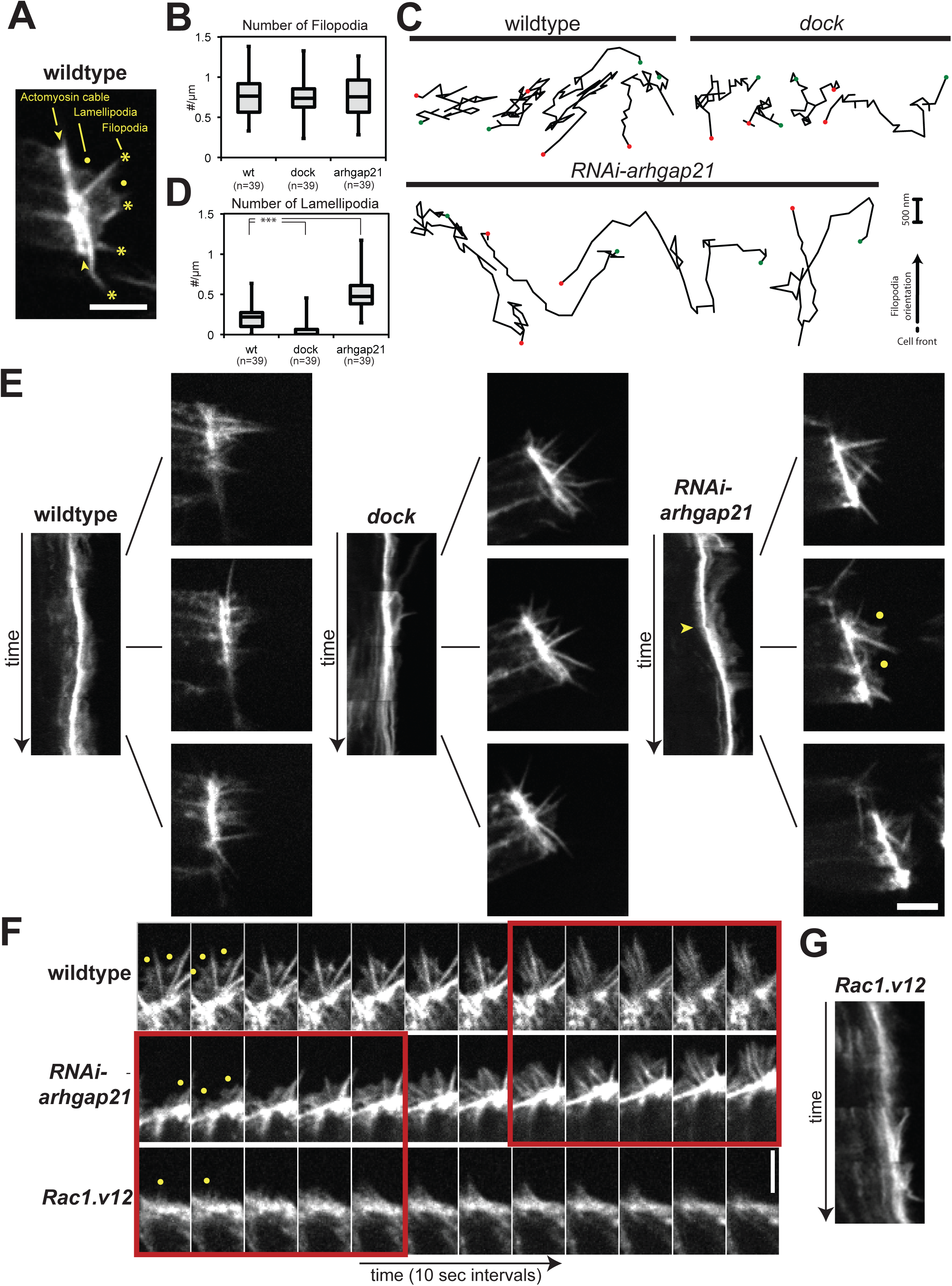
Dorsal closure leading edge actin structures. (A) Single plane of an epidermal leading edge (~0.1 μm) in en2.4-GAL4 UAS-LifeAct-GFP embryos. Actomyosin cable (yellow arrowhead), lamellipodia (yellow dot) and filopodia (yellow asterisk) are indicated. (B) Quantification of the number of filopodia detected per length of engrailed positive leading edge from still images in indicated embryo backgrounds. (C) Plots of filopodia tip position of three independent filopodia for each indicated genetic background. Green and red dots indicate first and last tip position. Each point represents a 2s interval. All tracks are oriented perpendicular to the leading edge. (D) Quantification of the number of lamellipodia detected per length of engrailed positive leading edge from still images in indicated embryo backgrounds. (E) For each indicated genetic background: (Left) Kymograph of the leading edge from Supplemental Movie S5. (Right) Still images that show leading edge state at indicated time point of the kymograph. For arhgap21 mutants: yellow arrowhead indicates start of a lamellipodia phase and dots indicate lamellipodia structures. Scale bar = 5 μm. (F) Montages of leading edge actin structures in indicated genetic backgrounds. Yellow dots indicate lamellipodia structures. Red boxes highlight biphasic similarities of arhgap21 mutants with wt and hyperactive Rac mutants. Scale bar = 5 μm. (G) Kymograph of the leading edge from Supplemental Movie S5.

Quantification of lamellipodia number revealed that a few lamellipodia occurred at the leading edge in the wt, while *dock* mutants rarely had any detectable lamellipodia; this is consistent with the classical lamellipodia activities of the elmo-dock complex (Figure 6D). In contrast arhgap21-depleted cells had significantly increased numbers of lamellipodia compared to wt (Figure 6D). In addition, arhgap21-depleted cell fronts appeared biphasic and transitioned between dominant filopodial and lamellipodial states compared to wt and *dock* mutants (Figure 6D and E, and Supplemental Movie S5). In arhgap21-depleted cells, when lamellipodia phases were synchronized within an engrailed positive stripe, a coordinated increase in leading edge movements was observed (Figure 6E and Supplemental Movie S5). Similar bursts in cell front progression were not detected in wt and *dock* mutants (Figure 6E).

Rac activity is known to promote lamellipodia formation (Guilluy et al., 2011). Therefore, these results suggest that a transient abnormal activation of Rac may occur in arhgap21-depleted cells. We tested the effects of Rac hyperactivation on the epidermal leading edge by expressing a constitutively active Rac (UAS-Rac1.V12) in engrailed stripes. Expression of Rac1.V12 often resulted in severe disruption of epidermal leading edges (data not shown). However engrailed-positive stripes, which maintained a well-defined leading edge, were dominated by lamellipodia and few filopodia (Figure 6F and Supplemental Movie S5). Furthermore, Rac1.V12-positive, lamellipodia-enriched cell fronts displayed a faster progression compared to wt fronts, similar to increased rate in arhgap21-depleted lamellipodial states (Figure 6E and G). Taken together these data imply that when the putative Rho GAP arhgap21 is depleted, the leading epidermal cells display aberrant rounds of Rac activity and inactivity (Figure 6F, red boxes).

### Arhgap21 regulates Rac and Rho1 activities in the epidermis

Rac activity was directly analyzed using the Parkhurst Pak1- and Pak3-based biosensors (Verboon and Parkhurst, 2015). The Pak1 biosensor gave no detectable signal during dorsal closure (data not shown). The Pak3 biosensor resulted in faint, ubiquitous membrane localization, and 80% of Pak3 biosensor homozygous embryos (15 seams from 10 embryos) showed the formation of sparse puncta in the epidermis leading edge prior to and after cadherin contact formation (Figure 7A and B). Pak3 biosensor puncta also appeared in the amnioserosa, with an apparent enrichment occurring during cell extrusion events (Figure 7C), which was not examined further. Engrailed expression of dominant negative UAS-Rac1.N17 was consistent with a Rac dependency of the puncta in the epidermis, but the sparse nature, limited localization, and 80% occurrence prevented conclusive determination. However, the ubiquitous, weak membrane localization of the Pak3 biosensor was not affected segmentally by engrailed-driven dominant negative Rac (data not shown). Therefore, we examined puncta formation in mutants of the elmo-dock complex, an established Rac regulator. In *elmo* mutants, Pak3-biosensor puncta were never detected (17 seams from 20 embryos), but membrane localization persisted (Figure 7A and B). These results suggests that the faint membrane localization was likely not specific for activated Rac, but the Pak3 biosensor puncta accurately reported Rac activity, and indicated that Rac activity in the epidermis is predominately focused prior to and immediately following cadherin contact formation.

**Figure 7.**
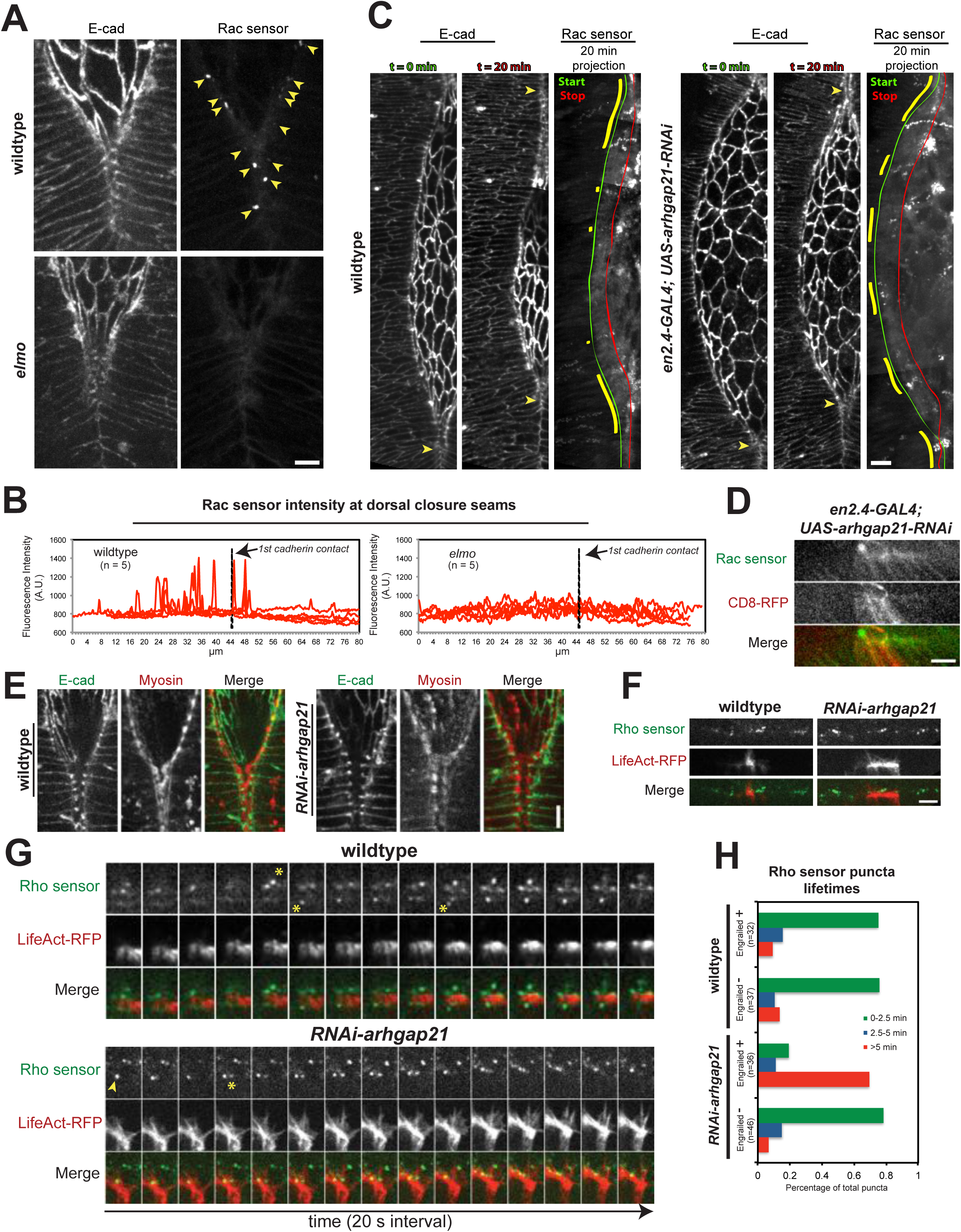
Arhgap21 regulates Rac1, Rho1 and myosin activities. (A) Maximal intensity Z-stack projections of wt and *elmo* mutant leading edges and cadherin seams, which express endogenous DE-cadherin-Tomato and spaghetti squash-driven Pak3-biosensor. Scale bar = 5 μm. (B) Each graph shows plots of absolute fluorescence intensities that were measured along a 20 pixel-wide line, which ran along and straddled one epidermal leading edge into cell-cell contact regions from 5 different embryos. Measurements were aligned to the newest cadherin. (C) Reassembled dorsal closure images from selected Z-planes of cadherin-labeled epidermal leading edges and seams that remove underlying amnioserosa planes. Embryos are engrailed-Gal4 and express endogenous DE-cadherin-Tomato and spaghetti squash-driven Pak3-biosensor with or without UAS-arhgap21-RNAi. For each set panels are (left) initial cell front, (middle) cell front after 20 minutes (right) Projection of maximal Pak3 localization over 20 minutes (Green = start, Red = stop of cell front). Yellow lines highlight regions of Pak3-biosensor puncta. Yellow arrows mark cadherin contact seams. Scale bar = 10 μm. (D) Maximal intensity projection (0.5 μm) of UAS-mCD8-mRFP; engrailed-Gal4; UAS-arhgap21 leading edge with squash-driven Pak3-GFP Rac sensor over 10 seconds. Orientation is cell front up. Scale bar = 5 μm. (E) Maximal intensity Z-stack projections of wt and ectodermally-expressed arhgap21 RNAi leading edges, which express endogenous DE-cadherin-GFP and exogenous spaghetti squash-driven spaghetti squash-mCherry. Scale bar = 5 μm. (F) Single focal plane (~0.1 μm) of wt and UAS-arhgap21 leading edges in an engrailed-Gal4, UAS-LifeAct-RFP background that express ubiquitin-driven Rho sensor-GFP. Orientation is cell front up. Scale bar = 5 μm. (G) Montages of single focal plane of wt and UAS-arhgap21 leading edges in an engrailed-Gal4, UAS-LifeAct-RFP background that express ubiquitin-driven Rho sensor-GFP. Yellow arrowhead shows foci with a > 5 min lifetime. Yellow asterisk indicates foci with a < 2.5 min lifetime. (H) Quantification of leading edge Rhosensor foci lifetimes obtained from movies collected at 2 s/frame

To examine the impact of arhgap21 on Rac activity, the formation of Pak3 biosensor puncta in the leading edge was monitored upon engrailed expression of arhgap21-RNAi. Analysis over 20 minutes that used DE-cadherin signal to select epidermal-specific Z-planes and exclude underlying amnioserosa biosensor signal revealed where Pak3 biosensor puncta formed along the closure. As expected, control cells mainly had activity near epidermal seams (Figure 7C). In arhgap21-depleted embryos, Pak3 biosensor puncta were frequently observed at the epidermis leading edge far away from the dorsal closure seams and these puncta occurred in segmental regions along the epidermis (Figure 7C). Co-expression of mCD8.mRFP and arhgap21-RNAi in engrailed stripes showed that the appearance of new Rac biosensor puncta at the leading edge (>25 cells away from the nearest canthi) occur 90% of the time (n=52 puncta from 3 embryos) in engrailed positive regions (Figure 7D). These results indicate that Rac activity is abnormally increased upon arhgap21 depletion, which is consistent with the transient formation of lamellipodia observed in arhgap21 mutants (Figure 6E).

Mammalian arhgap21 has Rho specificity in cells and has virtually no biochemical activity on Rac (Barcellos et al., 2013; Dubois et al., 2005; Lazarini et al., 2013). Therefore, the distribution of nonmuscle myosin II and the anillin Rho sensor were analyzed to assess Rho activity in arhgap21 mutants during dorsal closure. In wt cells, myosin II was enriched at the leading edge (Figure 7E). In arhgap21-depleted cells myosin also localized at the leading edge, but the accumulation was less even and compact compared to wt (Figure 7E) consistent with actin cable defects and tensions (Figure 5A and 4H). Embryos that expressed the anillin Rho sensor ubiquitously and UAS-LifeAct-RFP in engrailed stripes were analyzed in the presence and absence of UAS-arhgap21-RNAi. Little difference in overall Rho sensor levels was observed at the leading edge of engrailed-positive and -negative regions of wt and arhgap21-depleted embryos (Figure 7F). The lifetimes of Rho sensor puncta were analyzed and binned for all individual foci that appeared, disappeared or persisted during 5 min of image capture. This analysis revealed a population of stable Rho sensor puncta that occurred in engrailed-positive regions that were depleted for arhgap21 (Figure 7G and H). Together these data indicate that arhgap21 plays a role in Rac, Rho1 and myosin II regulation at the leading edge of epidermal cells during dorsal closure.

### The elmo-dock complex and arhgap21 regulate Rho1 activity independently

Since the elmo-dock complex inhibits Rho1 (Figure 3) and the GAP activity of arhgap21 is predicted to stimulate Rho1-GTP hydrolysis (Barcellos et al., 2013; Lazarini et al., 2013), a possible explanation for the synthetic defect between the two proteins (Supplemental Figure S1) is that Rho1-GTP is hyper-stabilized when both regulators are lost. To overcome previous attempts to analyze loss-of-function of both Rho family GTPase regulators, arhgap21 RNAi was expressed in engrailed stripes in *dock* mutant embryos. Compared to wt, these embryos displayed a severe disruption of the dorsal closure process, which included altered DE-cadherin signal and highly disorganized epidermal cells (Figure 8). Similar perturbations were observed in engrailed-positive (LifeAct-GFP) and engrailed-negative regions (Figure 8), which suggests possible non-cell autonomous defects. However, an unexpected early disruption of the amnioserosa layer prevented specific analysis of the epidermal leading edge structures in engrailed positive segments.

**Figure 8.**
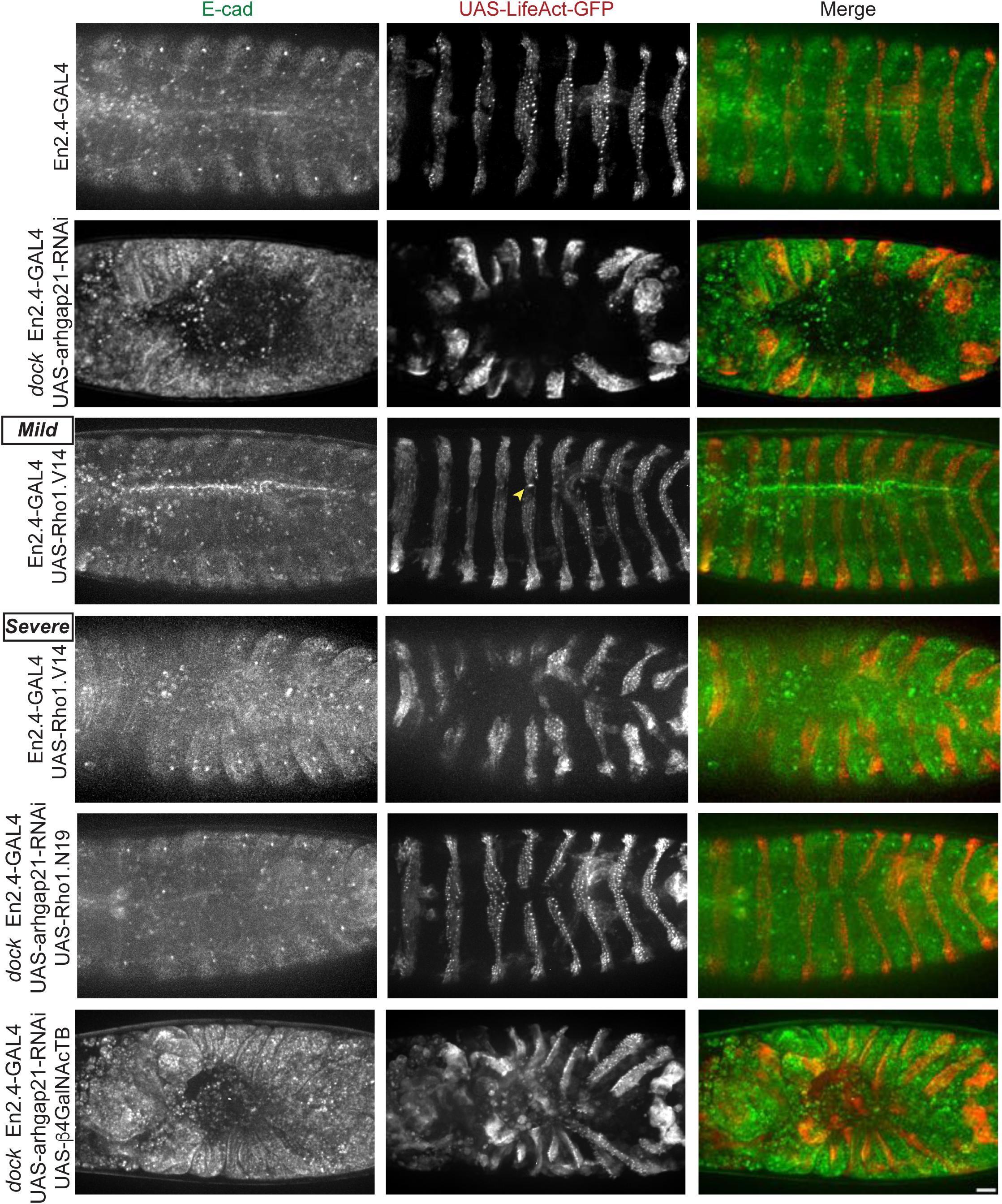
Elmo-dock complex and arhgap21 independently regulate Rho1. Maximal intensity Z-stack projections of embryos, which express endogenous DE-cadherin-Tomato and engrailed-Gal4, UAS-Lifeact-GFP in indicated genetic backgrounds. Yellow arrowhead indicates a constricted cell front. Embryos were staged for appearence of epidermal denticles. Scale bar = 25 μm.

To analyze the effects of Rho1 hyperactivation, constitutively active Rho1 (UAS-Rho1.V14) was expressed in engrailed stripes. A gradient of perturbations were observed: 54% (n=24) of embryos displayed mild effects in dorsal closure and had a constriction of engrailed-positive leading edges (Figure 8), as previously described (Jacinto et al., 2002); 46% (n=24) of embryos had severe defects that included highly disorganized epidermal cells and an early collapse of amnioserosa cells (Figure 8). These data indicate that *dock*, arhgap21-depleted mutants phenocopy the most severe aspects of Rho1 hyperactivation in the epidermis during dorsal closure.

To test the possibility of Rho1 hyperactivation upon loss of dock and arhgap21 function a dominant negative Rho1 mutant (UAS-Rho1.N19) was co-expressed in the double mutants. Expression of Rho1.N19 resulted in embryos that displayed a far more mild dorsal closure phenotype with incomplete closures (Figure 8), which were reminiscent of earlier reports that analyzed Rho family GTPase mutants (Harden et al., 1999). This result revealed a partial suppression of *dock* arhgap21-depleted mutants with dominant negative Rho. The use of UAS-β4GalNAcTB, an unrelated secretory pathway processing protein, was not able to rescue *dock* arhgap21-depleted dorsal closure defects (Figure 8) and showed that the rescue was not a result of Gal4 dilution effects. These results indicate that both the elmo-dock complex and arhgap21 regulate Rho1 activity through independent pathways.

## DISCUSSION

The novelty of this study is that it defines precisely in time and space Rac and Rho activities that occur during dorsal closure at the epidermal leading edge and newly formed cadherin contacts. Moreover, it identifies roles for the atypical Rac GEF elmo-dock complex and the Rho GAP arhgap21 in the coordination of these Rac and Rho cycles during *in vivo* MET-like transitions of *Drosophila* (Figure 9A). Mutants of the elmo-dock complex resulted in a dorsal closure similar to wt until a very late stage. Despite these initial similarities the elmo-dock complex mutants were defective in the number of epidermal lamellipodia present at the leading edge (Figure 6), which suggests that lamellipodia are largely dispensable for the majority of the dorsal closure process. These results are consistent with current models that suggest that the amnioserosa reduction provides the bulk force that drives dorsal closure while epidermal cell migratory contributions are minimal (Blanchard et al., 2010; Pasakarnis et al., 2016; Wells et al., 2014). In addition, Rho1, myosin II and tension regulation at new epidermal cadherin contacts were perturbed in elmo-dock complex mutants (Figure 2 and 3). Dual roles for the elmo-dock complex have been found in mammalian cells at lamellipodia and initial cell-cell contacts (Côté and Vuori, 2007; Toret et al., 2014a). The conservation of activities reveals that despite the differences between mammalian wound healing and dorsal closure systems, the elmo-dock activities identified in *Drosophila* likely generally apply widely to MET-like processes.

**Figure 9.**
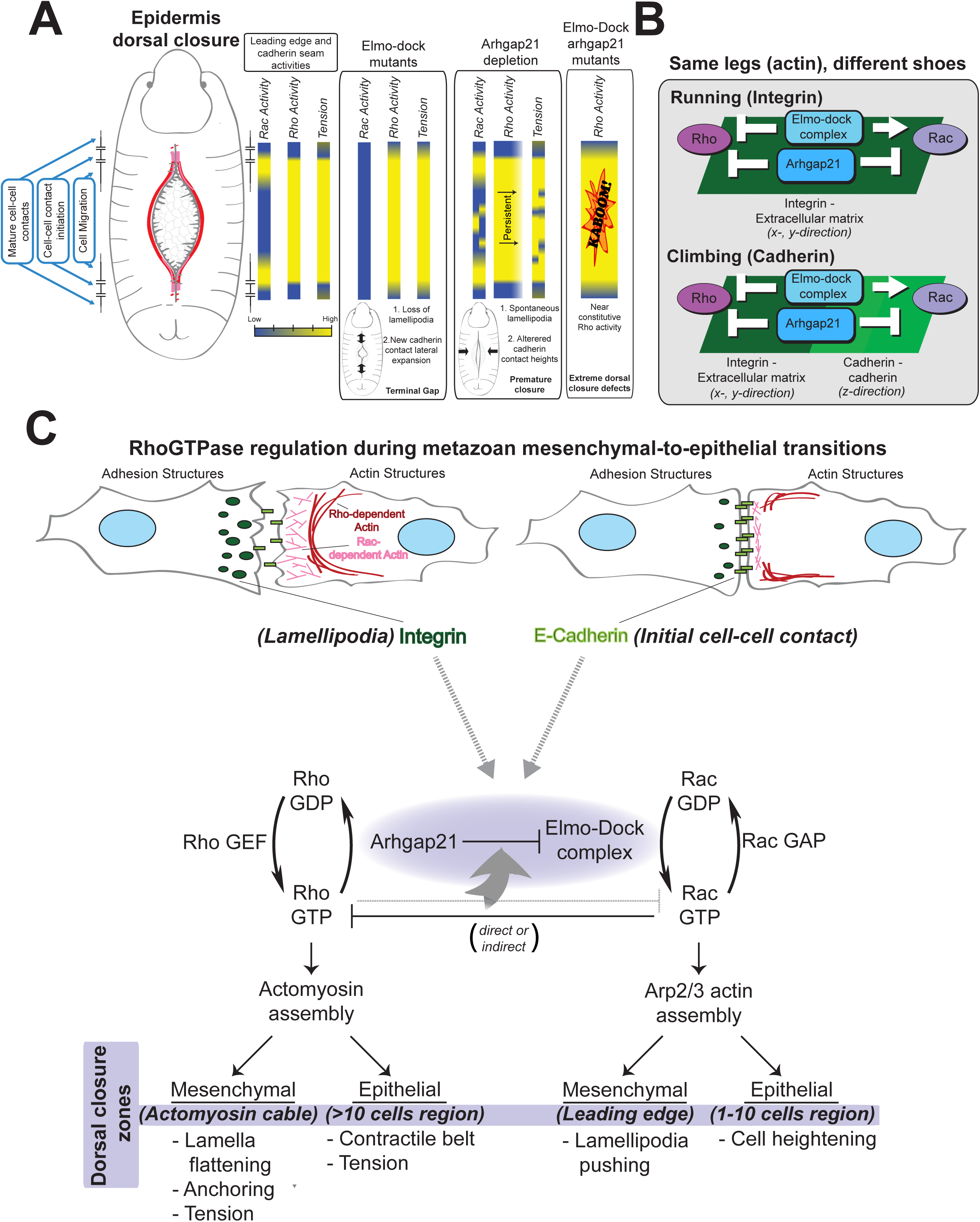
Model of elmo-dock complex and arhgap21 function during MET. (A) Schematic of the relative leading and cell-cell contact states during epidermis dorsal closure and heat maps that approximate Rac, Rho, and tension activities in wt and mutants. (B) Schematic showing parallel activities during cell migration and initial cell-cell contact formation. (C) A genetic pathway for Rho GTPase cycles during MET.

The elmo-dock complex drives Rac activation and actin rearrangements upon E-cadherin engagement in mammalian cells, but the role and consequences of actin reorganization at new cell-cell contacts could not be addressed in mammalian cells as cadherin contacts collapse upon elmo-dock depletion (Toret et al., 2014a). Elmo-dock complex-mediated Rac activation in lamellipodia is essential for membrane extension (Côté and Vuori, 2007), and an analogous role at new cadherin contacts would drive contact heightening by pushing membranes in the Z-direction up the cadherin substrate provided by the adjacent cell (Figure 9B). A membrane extension role can account for the cell height defect observed at new DE-cadherin contacts during dorsal closure in *elmo-dock* mutants, and may explain why initial E-cadherin contacts collapse in MDCK cells depleted for elmo2 (Toret et al., 2014a).

This study identifies a new, critical physical state of late dorsal closure (post-zippering/canthus) after epidermal DE-cadherin contacts form. In this region in wt embryos, a loss of tension at new DE-cadherin contacts is coordinated with a decrease in Rho activity and myosin localization. This region presents an unexpected, novel cadherin contact that experiences little to no tension and has major implications for force-dependent protein interactions that occur at these junctions, such as vinculin (Hoffman and Yap, 2015; Lecuit and Yap, 2015). It remains to be identified how these junctions remain unaffected by the forces of the adjacent actomyosin cable and mature contacts. The formation of this tension free zone has a major impact on the normal embryo development and its absence results in a lateral expansion of new contacts. As a result of this lateral expansion, as new contacts form in *elmo-dock* mutants, the epidermis becomes elongated, which results in a leading edge that is progressively squeezed. As dorsal closure proceeds, this effect creates the terminal gaps that occur in *elmo-dock* mutants (Figure 9A).

In *Drosophila*, depletion of arhgap21 resulted in embryos that complete epidermal closure faster than wt (Figure 9A). Arhgap21-depleted embryos displayed complex epidermal cell phenotypes (a fragmented actomyosin cable, bimodal leading edge tensions, transient Rac and lamellipodia states and shorter and taller cadherin contacts). The fragmented actomyosin cable can explain the observed binary tensions. Curiously, the actomyosin cable and tension behaviors at the leading edge mirrored more closely Rac than Rho biosensor behavior upon loss of arhgap21, which hints at a myosin and Rac link (Figure 9A). The increase in lamellipodia protrusions can explain the faster migration of the epidermis (Figure 9A and B). In wt embryos, the speed of the epidermal leading edge and the reduction of the amnioserosa were equal, which suggests that these two processes are coupled to maintain cell migratory contributions at a negligible level, whereas in *arhgap21* mutants they were decoupled. However, it remains unclear how amnioserosa contraction and epidermal cell migration pathways crosstalk to coordinate these two closure rates. Unexpectedly, lateral filopodial dynamics were decreased in *elmo-dock* mutants and increased in *arhgap21* mutants. This requires further study, but could be due to indirect consequences of dynamic lamellipodia changes that affect the associated filopodia or to unexplored links with Cdc42. During dorsal closure, arhgap21 mutants also had defects in new cadherin contact cell heights and tensions. A transient regulation of Rac activity at new contacts, similar to arhgap21-depeleted leading edges, could account for under- and over-heightened cadherin contacts observed upon loss of arhgap21 (Figure 9B). Depletion of mammalian arhgap21 results in faster cell migration rates, although the cellular consequences of arhgap21 depletion were not defined (Barcellos et al., 2013; Bigarella et al., 2009; Lazarini et al., 2013). Additionally, mammalian arhgap21 has an undefined role at cadherin contacts and was reported to localize at new E-cadherin contacts with kinetics that resemble the Elmo2-Dock1 complex (Barcellos et al., 2013; Sousa et al., 2005; Toret et al., 2014a). Together these results suggest that, similar to the elmo-dock complex, arhgap21 likely also has conserved roles in invertebrate and mammalian MET-related processes. In mammalian cells, arhgap21 was first reported to activate predominately Cdc42 *in vitro*, but later mammalian tissue culture studies favored a role in the activation of RhoA and RhoC, over Cdc42 (Barcellos et al., 2013; Dubois et al., 2005; Lazarini et al., 2013). Surprisingly, the *in vivo* dorsal closure phenotype (faster closure) and Rac biosensor data favor a potential transient Rac GAP role for arhgap21. In contrast, Arhgap21 depletion also stabilized Rho sensor foci, which also supports a conserved Rho GAP function (Figure 9A and B). Curiously, the mammalian studies that identified Rho activation by arhgap21 also report an increase in cell migration rates suggesting that this Rac effect may also be conserved (Lazarini et al., 2013). Moreover, a synthetic lethal genetic interaction occurs upon loss-of-function of both arhgap21 and the elmo-dock complex that mirrors the severest defects caused by constitutively active Rho1 expression and is suppressed by dominant negative Rho1. This agrees with the possibility that both proteins function in dual pathways that inactivate Rho1 at the leading edge and new contacts of epidermal cells (Figure 9B). A precise understanding of this complex hyperactive Rho1 dorsal closure defect (Figure 9A) requires further study.

The complex and confounding regulation of Rac and Rho identified in this study and previous mammalian works can be accounted for in a model in which the classical inhibitory relationship between Rho and Rac (Burridge and Wennerberg, 2004) is transferred to the Rho GAP and Rac GEF (Figure 9C). The removal arhgap21 results in an inability to stimulate Rho1 GTPase activity directly (persistent Rho activation), and also a failure to inhibit elmo-dock-mediated Rac activation. Improperly regulated and transient Rac activity would still directly or indirectly inhibit Rho processes like actomyosin-generated tension. For example, high Rac activity would result in loss of actomyosin cable/ tension (Figure 9A). Loss of elmo-dock complex function would prevent Rac activity, and thus not inhibit Rho until the Rho GAP, or other secondary mechanisms could compensate. This would explain the tapering off of Rho, Myosin, and tension levels at new cadherin contacts in elmo-dock mutants. Loss of both elmo-dock and arhgap21 would prevent Rac activation, but also all Rho inactivation (Figure 9A). The mechanisms that underlie arhgap21 regulation of the elmo-dock complex may be direct or indirect, and future analysis of biochemical activities, spatio-temporal protein dynamics and associated scaffolding proteins are necessary to resolve this relationship.

This study identifies for the first time a new GEF-GAP protein partnership as a major regulator of the inverse relationship of Rac and Rho cycles specifically during leading edge to cadherin contact transitions. Curiously, Arhgap21 was identified as an EMT protein (Barcellos et al., 2013), but it remains unclear how the MET-related functions identified here relate to the disruption of cadherin contacts and establishment of a leading edge. Other GEF and GAP proteins known to function at cadherin sites may act during non-MET-related processes such as at mature junctions or during other cadherin contact expansions (e.g.) after cytokinesis or T1-T3 transitions (McCormack et al., 2013). We identify striking parallels between *Drosophila* and mammalian systems despite the apparent mechanistic differences between the processes of dorsal closure, wound healing and cell pairs forming cadherin contacts. Therefore, dorsal closure is a powerful model to study MET-related processes and these findings will likely translate broadly across metazoans. Rho regulators studies on mammalian neural tube closure, palatogenesis and myogenesis are necessary to further define common features.

## ACKNOWLEDGEMENTS

We thank WJN, EB, QM, and AGDLB for critical reading of this manuscript. This project was supported by la Fondation ARC. Dossier n° PDF20131200557, CNRS and Aix-Marseille Univ, the labex INFORM (grant ANR-11-LABX-0054). We thank the IBDM imaging facility for imaging support and acknowledge France-BioImaging infrastructure supported by the Agence Nationale de la Recherche (ANR-10-INSB-04-01, call “Grand Emprunt”). The Le Bivic group is an “Equipe labellisée 2008 de La Ligue Nationale contre le Cancer”. We thank the TRiP at Harvard Medical School (NIH/NIGMS R01-GM084947) for providing transgenic RhoGAP19D-RNAi fly stock.

## MATERIALS AND METHOD

### Drosophila

All *Drosophila* work was carried out at 25°C. Fly strains used in this study are listed in Supplemental Table S1. To visualize dorsal closure embryos were collected 12–14 hr after egg laying, dechorionated for 1 min in bleach, aligned and mounted on a coverslip in Halocarbon 200 oil (25073, Polysciences, Inc).

### Microscopy

Embryos were imaged on a Nikon Ti-E inverted microscope equipped with a Yokogawa CSU-X1 spinning disk and an EM-CCD Camera (Photometrics Evolve 512). A 20× air 40× 1.25 N.A. water-immersion and 100× 1.4 N.A. oil-immersion objectives were used. Image acquisition was performed with MetaMorph. For 20× or 40× images, 60 planes (1.2 or 0.2 μm, repsectively) were captured that spanned the dorsal side of the embryo. For 100× images 60 planes (0.1 μm) were captured that spanned the epidermis leading edge or dorsal closure seam. GFP and RFP channels were captured sequentially. Laser power kept constant between experiments.

Laser ablations were performed on 1W average power, femtosecond, near infrared laser equipped with spinning disc microscope for imaging. To ablate cell-cell contacts, point ablations were done with an average power of 350-450mW (sample dependent) and the duration of the exposure was 200ms.

### Image Analysis

All images were analyzed with ImageJ (http://rsb.info.nih.gov/ij/). All movie rates are listed with corresponding Supplemental Movie descriptions.

Amnioserosa extrusion rates were calculated by imaging early stage 15 embryos for one hour (1min/ frame). The number of extruding cells (a shrinking, collapse, and loss of a DE-cadherin positive cells) over the course of the movie was divided by the total number of initial amnioserosa cells.

Actomyosin pulse quantifications were calculated from the time of appearance to the disappearance of a medial population of LifeAct-RFP in amnioserosa cells during mid/late Stage 15. Data was collected from 30 sec/frame movies.

Laser ablation analysis were done by measuring the distance between the vertices of the ablated cell-cell contact over the time. A linear fit for first 4 points were used to the graph the measured distance over time, which gave the value of initial velocity value.

Closure rates were calculated by measuring the change in linear distance that separates the cadherin-labeled leading edge or LifeAct-labeled aminoserosa borders during dorsal closure over 5 min intervals. Thirty to forty measurements were pooled from three to four embryos for each analysis. Data was collected from 1 min/frame movies.

Filopodia and lamellipodia counts were obtained from single plane, still images captured as seen in Figure 5A. The numbers of filopodia or lammellipodia webs were counted and divided by the length of the actin cable/segment length. For filopodia tip tracking. The x,y position of the filopodia tip was recorded at 2 s intervals and plotted on an x, y project. Green dot indicates the initial position and red dot indicates the last position recorded. Tracks were oriented so that the actin cable is down.

For all box and whisker plots, the ends of the box mark the upper and lower quartiles, the central horizontal line indicates the median, and the whiskers indicate the maximum and minimum values. For graphs (**) and (***) indicate P<0.05 and P<0.0001 (unpaired T-test), respectively.

## SUPPLEMENTAL FIGURE LEGENDS

**Supplemental Figure S1.**
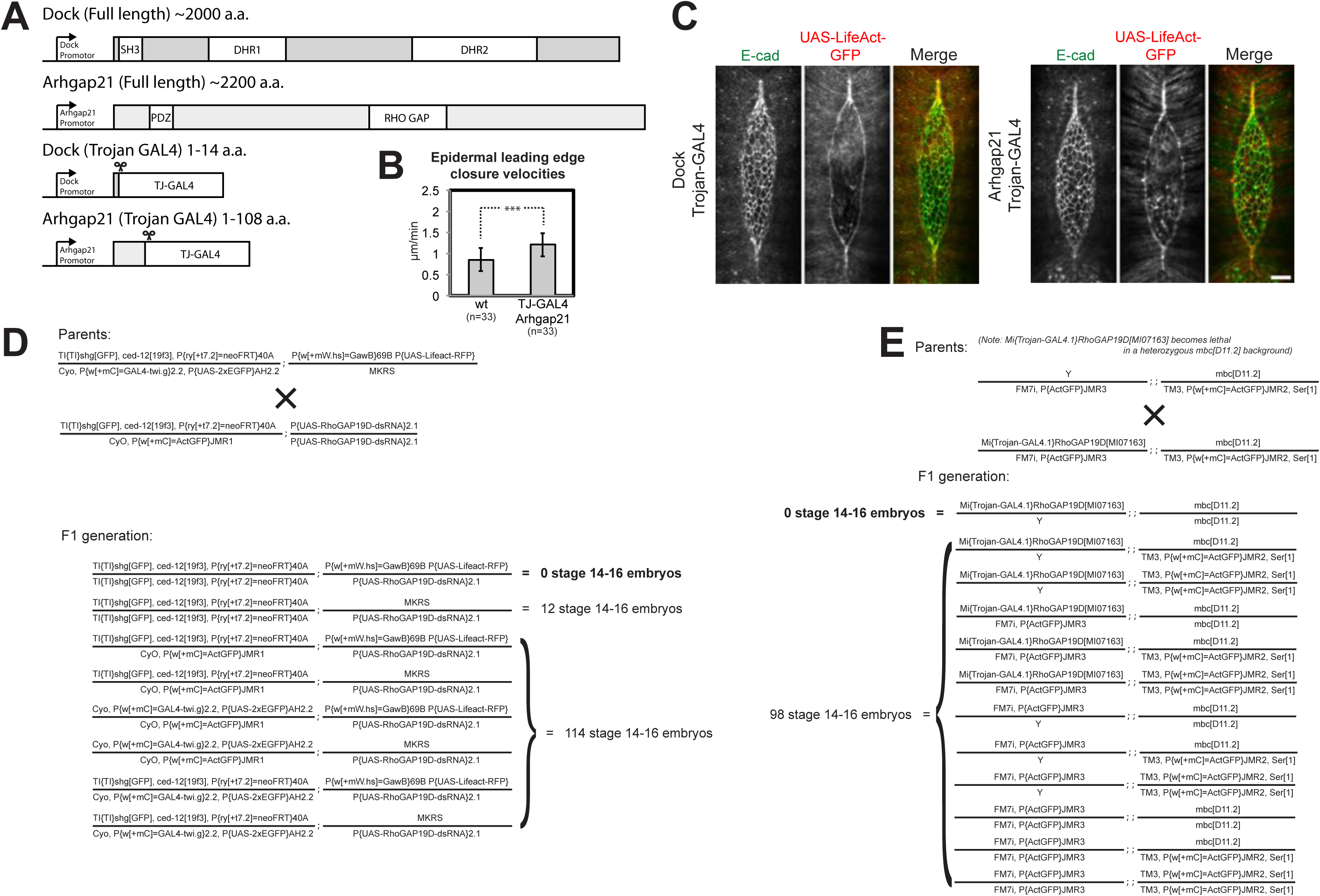
Elmo-dock and arghap21 supplemental results. (A) Schematic of Dock and Arhgap21 proteins and corresponding Trojan-GAL4 mutants. (B) Quantifications of the change in linear distance that separate leading edges (DE-cadherin-GFP) during dorsal closure in 5 min intervals. Measurements pooled from two embryos from each of the indicated conditions. (C) Maximal intensity Z-stack projections of entire late stage dorsal closure in of embryos that express endogenous DE-cadherin-Tomato and indicated Trojan GAL4 with UAS-LifeAct-GFP. Scale bar = 25 μm. (D) Genetic cross to obtain *elmo-dock* arghap21-depleted embryos and observed progeny counts (E). Genetic cross to obtain *elmo-dock arghap21* mutant embryos and observed progeny counts

## Supplemental Movie LEGENDS

**Supplemental Movie S1. Dorsal closure defects of *elmo* and *dock* mutants.** Maximal intensity Z-stack projections of the dorsal side of endogenous DE-cadherin-GFP-expressing wt (left), *elmo* (middle) and *dock* (right) mutant embryos (1 min/frame). Scale bar = 25 μm.

**Supplemental Movie S2. Dorsal closure in *elmo dock* mutants.** Maximal intensity Z-stack projections of the dorsal side of endogenous DE-cadherin-GFP-expressing *elmo dock* mutant embryos (1 min/frame). Scale bar = 25 μm.

**Supplemental Movie S3. Leading edge and cell-cell tensions.** Single focal plane of endogenous DE-cadherin-GFP-labeled contacts (250 ms/frame). Regions shown are wt (left panels) and *elmo* mutants (right panels) with old cell-cell contacts (top), new cell-cell contacts (middle) and leading edge (bottom). Scale bar = 5 μm.

**Supplemental Movie S4. Dorsal closure defects upon arhgap21 depletion.** Maximal intensity Z-stack projections of the dorsal side of wt (top) and UAS-arhgap21 (bottom) embryos. Embryos have endogenous DE-cadherin-GFP (green) and ectodermal Gal4 UAS-LifeAct-RFP (red) (1 min/frame). Scale bar = 25 μm.

**Supplemental Movie S5. Lamellipodia and filopodia dynamics during dorsal closure.** Single planes of epidermal leading edge in en2.4-GAL4 UAS-LifeAct-GFP embryos (2 s/frame). Backgrounds shown are wt (far left), *dock* (left), UAS-arhgap21-RNAi (right), UAS-rac.V12 (far right). Scale bar = 5 μm.

## SUPPLEMENTAL TABLE LEGEND

**Supplemental Table S1.**
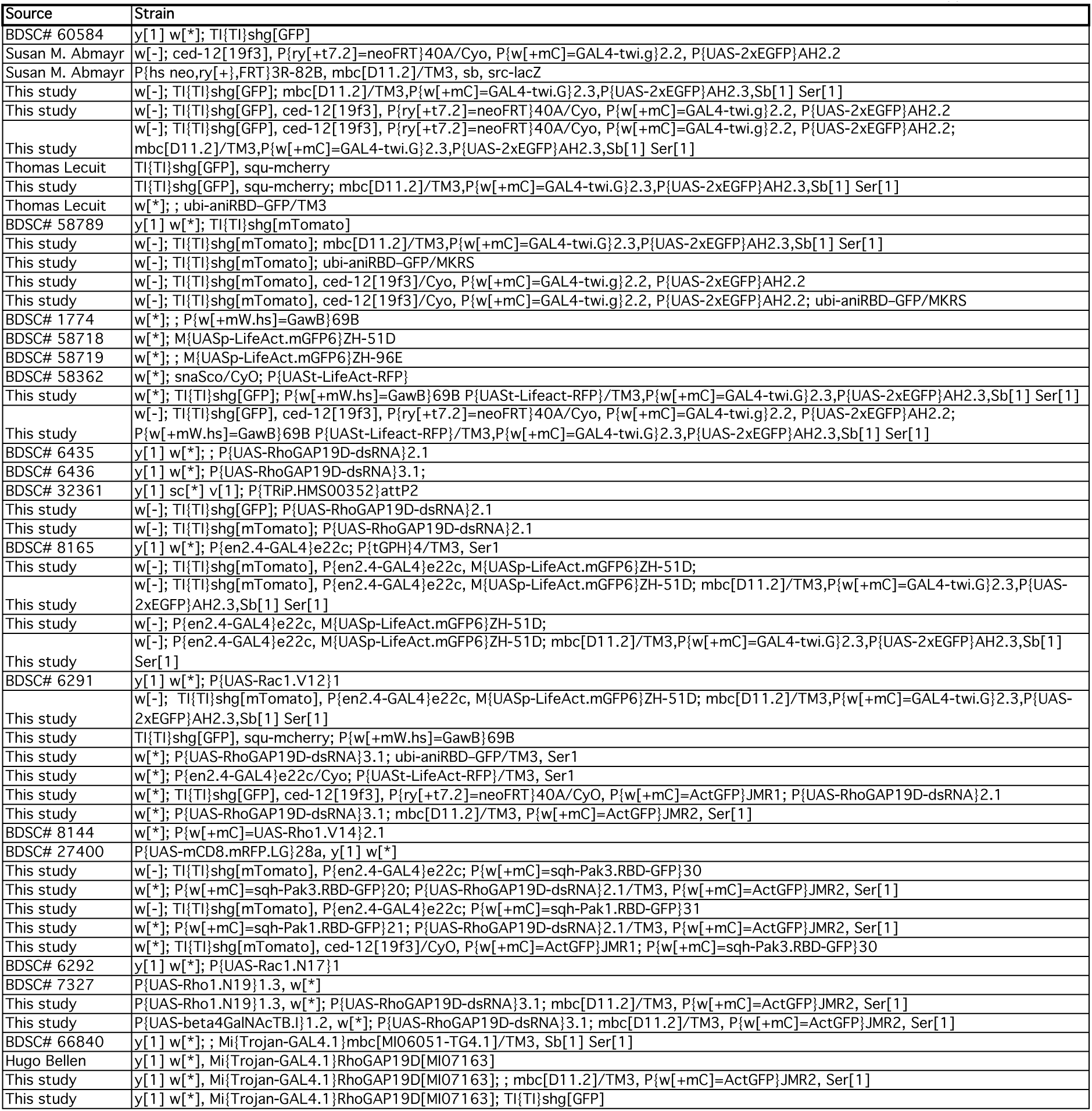
*Drosophila* strains. *Drosophila* used in this study

